# An explicit formula for a dispersal kernel in a patchy landscape

**DOI:** 10.1101/680256

**Authors:** Ali Beykzadeh, James Watmough

**Affiliations:** University of New Brunswick

## Abstract

Integrodifference equations (IDEs) are often used for discrete-time continuous-space models in mathematical biology. The model includes two stages: the reproduction stage, and the dispersal stage. The output of the model is the population density of a species for the next generation across the landscape, given the current population density. Most previous models for dispersal in a heterogeneous landscape approximate the landscape by a set of homogeneous patches, and allow for different demographic and dispersal rates within each patch. Some work has been done designing and analyzing models which also include a patch preference at the boundaries, which is commonly referred to as the degree of bias. Individuals dispersing across a patchy landscape can detect the changes in habitat at a neighborhood of a patch boundary, and as a result, they might change the direction of their movement if they are approaching a bad patch.

In our work, we derive a generalization of the classic Laplace kernel, which includes different dispersal rates in each patch as well as different degrees of bias at the patch boundaries. The simple Laplace kernel and the truncated Laplace kernel most often used in classical work appear as special cases of this general kernel. The form of this general kernel is the sum of two different terms: the classic truncated Laplace kernel within each patch, and a correction accounting for the bias at patch boundaries.

## 1 Introduction

Integrodifference equations are often used to model the distribution of annual species with distinct growth and dispersal stages across a landscape. The general form of an integrodifference equation (IDE) modelling growth and dispersal is as follows:

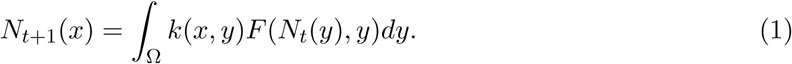

Here *N*_*t*_(*x*) is the population density at a point *x* ∈ Ω at the end of the dispersal period in year *t*, the growth function, *F*, is the density of dispersers, or new recruits following the growth phase of year *t*, and the dispersal kernel, *k*(*x, y*), is the probability a disperser originating at a point *y* ∈ Ω disperses to *x* [Kot and Schaffer, 1986]. Note that we do not assume all dispersers survive the dispersal phase. Hence, ∫_Ω_ *k*(*x, y*) *dy* ≤ 1. The integral operator in (1) allows us to accommodate a wide variety of empirically determined dispersal patterns.

IDEs with a discontinuous population density have been recently analyzed in patchy landscapes equipped with discontinuous kernels [see e.g., Neupane and Powell, 2015]. Many of these studies restrict the analysis to the case with no bias at patch boundaries [see e.g., Garlick et al., 2011]. However, the degree of bias has direct effect on, for example, the speed of spread [Lutscher and Musgrave, 2017], and should be included in dispersal models. There is an explicit formula for a discontinuous dispersal kernel in case of periodic infinite patches with considering the habitat preference [Musgrave and Lutscher, 2014]. Yurk and Cobbold [2018] presented a homogenization approach to the multi-scale problem of how individual behavioural responses to sharp transitions in landscape features, such as forest edges, affect population-dynamical outcomes. In their work, they treated the special case of a periodic environment consisting of two types of alternating patches, but the theory carries over to other periodic settings. We have obtained a general *m*-patch kernel including multiple patches with movement bias at the boundaries. The *m*-patch kernel extends several previous models for dispersal.

Many heterogeneous landscapes can be reasonably approximated by a mosaic of homogeneous patches, where individuals can respond to the boundaries by changing dispersal characteristics and by possibly immediately reversing direction to stay within the same patch. Musgrave [2013] showed that a certain formulation of a random walk with bias at the boundaries on such landscapes leads to discontinuities in the dispersal kernel at the patch boundaries. Dispersal bias, while not necessary for persistence, can significantly increase chances of persistence [Lutscher et al., 2010]. More generally, there are many behaviours, especially in two or three dimensions involving responses to boundaries and movement along boundaries.

Here our aim is to derive a general dispersal kernel for the most straightforward one-dimensional case. We first focus on a heterogeneous landscape that can be approximated by homogeneous patches, then we partition the heterogeneous landscape into pieces with homogeneous dynamics on each piece. Within each patch, dispersal follows a simple random walk with a constant step size, mortality and settling rates [as per Ovaskainen and Cornell, 2003]. We obtain an explicit formula for the kernel as piecewise exponential function with coefficients and rates determined by the inverse of a matrix of model parameters.

## 2 General dispersal kernel

### 2.1 Kernel Derivation

Let {*a*_0_, …, *a*_*m*_} denote the positions of the *m* +1 interface points of Ω = ℝ, and let Ω_*i*_ = (*a*_*i*−1_, *a*_*i*_), *i* ∈ {1, …, *m*} denote the bounded patches, which are accompanied by two semi infinite patches, Ω_0_ = (−∞, *a*_0_) and Ω_*m*+1_ = (*a*_*m*_, ∞). For each patch Ω_*i*_, the settling rate, the death rate, and the motility are denoted by *α*_*i*_, *β*_*i*_, and *ν*_*i*_, respectively. The degree of bias, −1 *< z*_*i*_ *<* 1, at each interface determines two probabilities, (1 − *z*_*i*_)*/*2 and (1 + *z*_*i*_)*/*2, of moving to the left and to the right at boundary *i*, respectively. Ovaskainen and Cornell [2003] showed that the dispersal kernel is the Green’s function, *k*(*x, y*), satisfying the partial differential equation

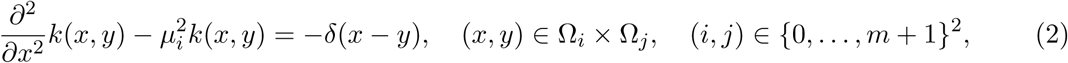

where 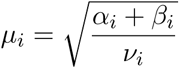, and the following conditions:

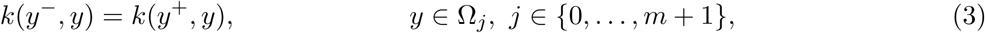

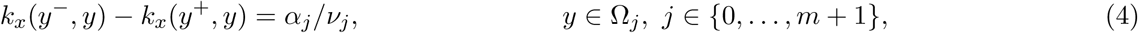

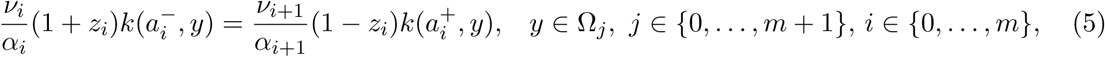

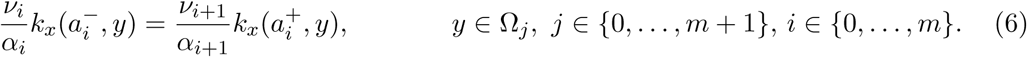

Condition (3) ensures the kernel is continuous at *x* = *y*; Condition (4) arises from a matching condition [see Keener, 2000]; Condition (5) is a jump condition; and finally, Condition (6) is the flux balance condition and represents continuity of the flux across the interface, so that no individuals are added or removed at the interface [Musgrave, 2013]. To ensure the kernel is bounded, we also add the conditions

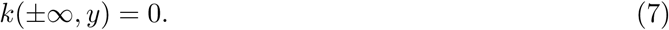

We expect the kernel to be piecewise continuous with possible discontinuities along the lines *x* = *a*_*i*_, *i* ∈ {0, …, *m*} and *y* = *a*_*j*_, *j* ∈ {0, …, *m*}. Let *k*_*ij*_ denote the portion of the kernel representing dispersal from patch Ω_*j*_ to patch Ω_*i*_. That is, *k*(*x, y*) = *k*_*ij*_(*x, y*), for (*x, y*) ∈ Ω_*i*_ × Ω_*j*_, (*i, j*) ∈ {0, …, *m* + 1}^2^.

Using *E*_*i*_ to denote the functions

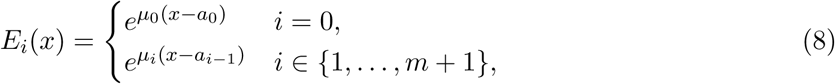

which are solutions to 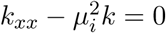, we can express each piece of the kernel in the form

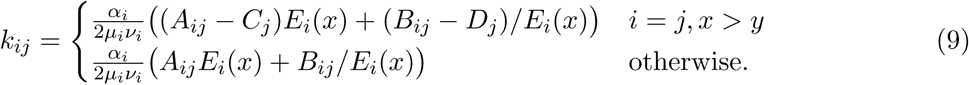

It follows immediately from Condition (7) that

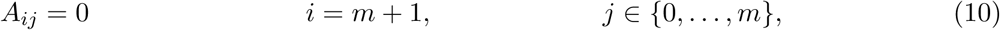

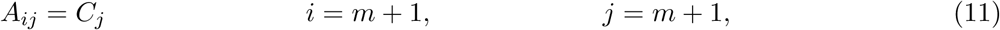

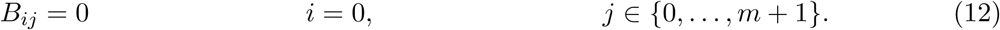

From continuity at *x* = *y* we have

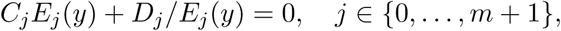

and from the matching condition on *k*_*x*_ at *x* = *y* we have

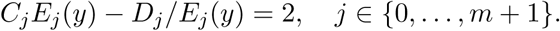

These can be solved for *C*_*j*_ and *D*_*j*_ to obtain

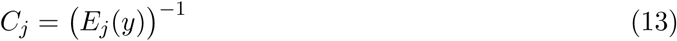

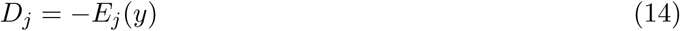

Upon substituting the expression for *C*_*m*+1_ into Equation (11), we find that

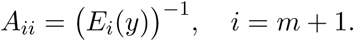

From Conditions (5) and (6), for (*i, j*) ∈ {0, …, *m*}×{0, …, *m*+1}, we have the interface conditions

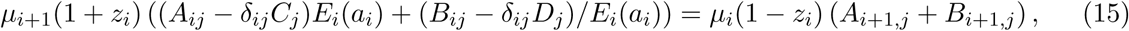

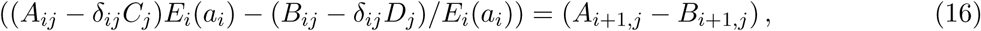

where *δ*_*ij*_ denotes the Dirac-delta.

Substituting Equations (13) and (14) into Equations (15) and (16) leads to the conditions

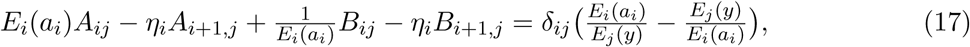

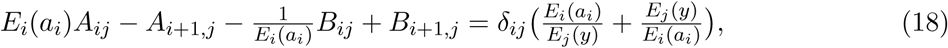

where

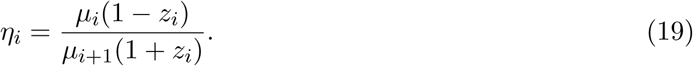

Equations (17) and (18) along with Equations (10)-(12) are a linear system of 2(*m* + 2)(*m* + 2) equations for the coefficients *A*_*ij*_ and *B*_*ij*_ with (*i, j*) ∈ {0, …, *m* + 1}^2^. Solving this system for the coefficients gives us a piecewise-defined form of the kernel as expressed by Equation (9)

### 2.2 Matrix Form of the Kernel

The system derived in the previous section is linear, implying that the kernel can be expressed in a compact matrix form. This result can be exploited to greatly reduce the computations required to numerically compute the kernel and emphasizes the similarity of the general kernel with the simpler Laplace kernels and truncated Laplace kernels appearing in the literature.

Let *ϵ*_*i*_ = *E*_*i*_(*a*_*i*_), and let *T* and *W* denote the following blocked matrices of parameters:

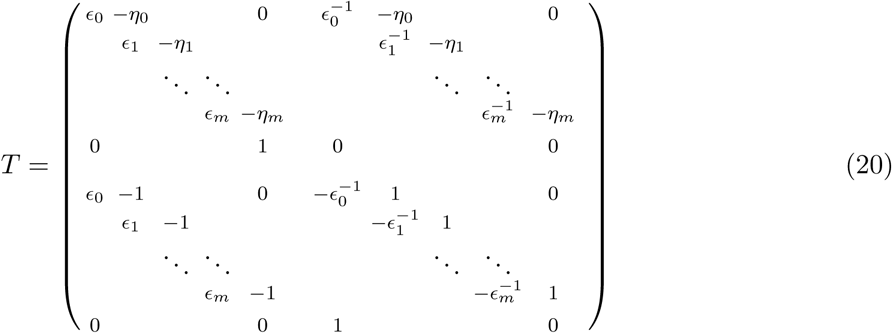

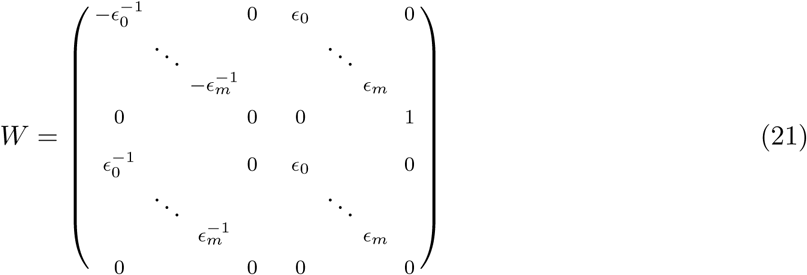

If we let *A* and *B* denote the two (*m* + 2) × (*m* + 2) matrices of coefficients, and we define

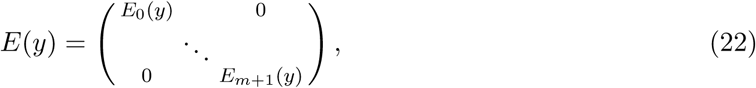

we can write the system of equations from the previous section as

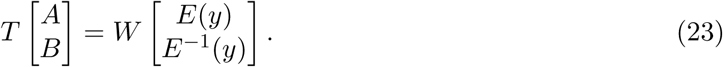

Consequently,

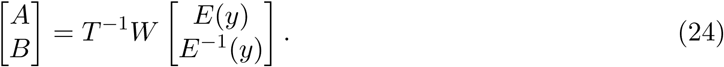

Thus Equation (9) has the matrix form

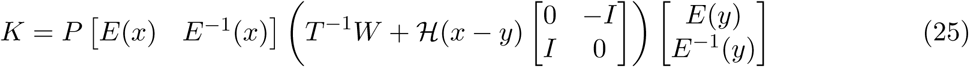

where *P* is a diagonal matrix with diagonal entries 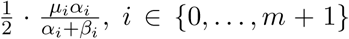, and ℋ is the Heaviside step function.

It is useful to rearrange Equation (25) to bring out a comparison between the general form of the kernel and the simple Laplace kernel. To this end, note that 1 − ℋ(*x* − *y*) = ℋ(*y* − *x*), and so

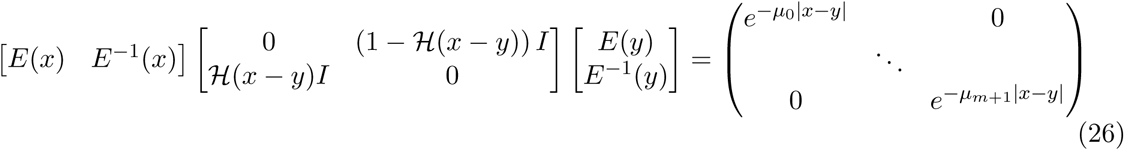

Hence,

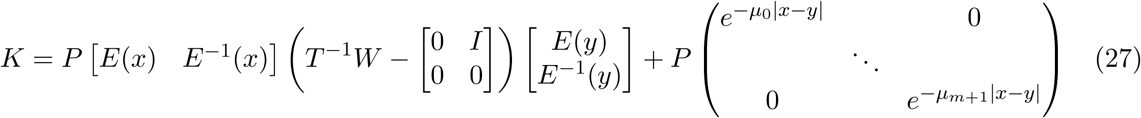

The reciprocal of *ϵ*_*i*_ can be interpreted as a measure of patch-width. Specifically, if we define the homogeneous Laplace kernel by

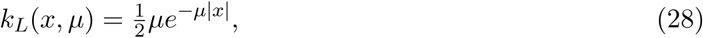

then for 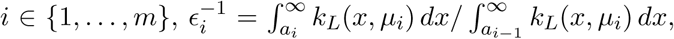, which is the probability a given individual settles beyond *a*_*i*_ given that they settle beyond *a*_*i*−1_ and can be viewed as the fraction of dispersers reaching *a*_*i*−1_ that continue across *a*_*i*_. Similarly, *η*_*i*_ can be viewed as a measure of the retention at the interface: *η*_*i*_ is the ratio of the rescaled kernel across the interface. Note, however, that retention also depends on the entries of *P*, or more accurately on the jumps in motility, *ν*, and settling, *α*, across the interface, and the value of *z*_*i*_.

Since the entries of *T*^−1^*W* depend in general on dispersal parameters in every patch through the two parameter groups, *η* and *ϵ*, the component of the kernel, *k*_*ij*_ representing settlement in patch *i* by dispersers originating in patch *j*, also depends in general on all dispersal parameters. However, it is important to note that the matrix form of the kernel implies that the functional form of each piece *k*_*ij*_ is a linear combination of the four functions

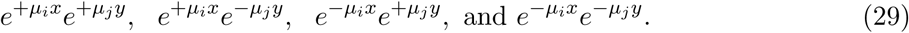

## 3 Special cases

### 3.1 Single interface

To illustrate the method, we first consider the simplest case where *m* = 0. That is, there is only a single interface point, with possibly different dispersal parameters on each side of the interface. From the definitions of *T* and *W*, and noting that *ϵ*_0_ = 1, we find that

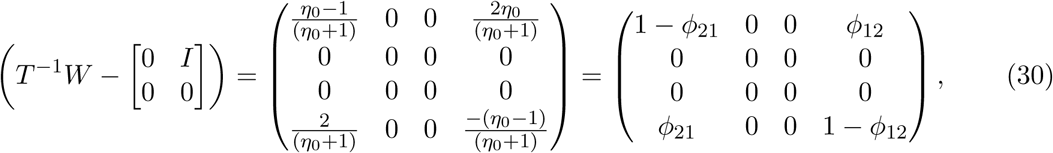

with 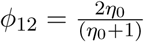 and 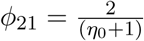. Hence, from Equation (27),

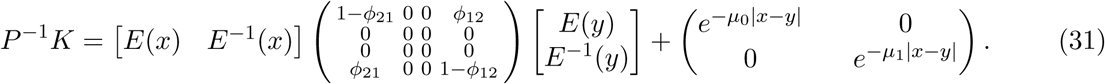

The *k*_*ij*_ entry of the kernel involves only the (*i* + 1)st row and (*j* + 1)st column of the product of the right hand side. For example, *k*_00_ involves the first row and first column of the product, which involves the first row of *E*(*x*) and *E*^−1^(*x*) and the first column of *E*(*y*) and *E*^−1^(*y*). Since *E*(*x*) is a diagonal matrix, this implies only the first and third rows and columns of *T*^−1^*W* are used in computing *k*_00_. Similarly, *k*_01_ uses only the first and third rows and the second and fourth columns of *T*^−1^*W*.

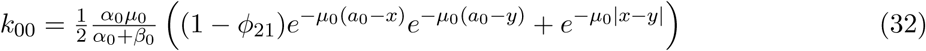

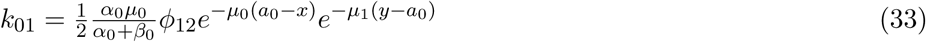

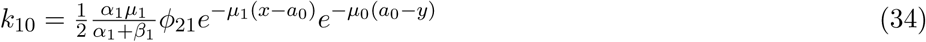

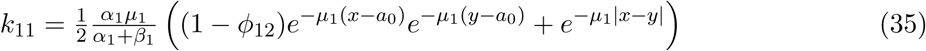

The pattern of zeros in *T*^−1^*W* after subtracting the identity from the upper right block arises from the condition that the kernel remain bounded. For a fixed *x*, when *y* → ∞ in Equation (33), the dispersal kernel goes to zero. We have the same limit for *k*, when *y* → −∞ in Equation (34).

For *m* = 0, we can, without loss of generality, assume *a*_0_ = 0. If we further assume that the habitat quality is identical in both patches (*µ* = *µ*_0_ = *µ*_1_, etc.), we have *η*_0_ = (1 − *z*_0_)*/*(1 + *z*_0_), and after dropping the unnecessary subscripts on the parameters the components of the kernel simplify to

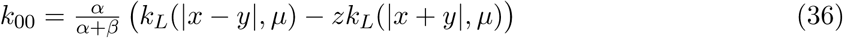

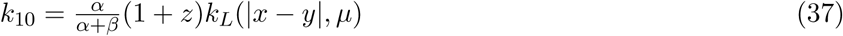

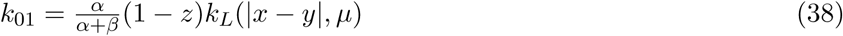

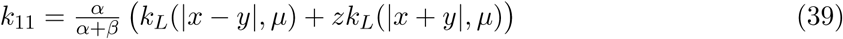

The first term, *k*_*L*_(|*x* − *y*|, *µ*) in *k*_00_ and *k*_11_ is the probability density for a disperser following a Homogeneous Laplace kernel dispersing a distance |*x* − *y*|, and the remaining term in each piece, *k*_*L*_(|*x* + *y*|, *µ*), is the probability density for the same disperser travelling a net distance |*x* + *y*|. This second distance is the net distance travelled by a disperser settling at the reflection of *x* across the interface. With no bias, *z* = 0, each piece of the kernel reduces to the homogeneous Laplace kernel. With *z >* 0, a fraction *z* of dispersers are reflected back across the interface, thus the terms of *k*_11_ are the following probabilities, with *x* and *y* positive: *k*_*L*_(|*x* − *y*|, *µ*) is the probability density for a disperser travelling from *y* to *x*; *k*_*L*_(|*x* + *y*|, *µ*) is the probability density for a disperser travelling from *y* to (−*x*) in absence of bias; with bias, a fraction *z* of these dispersers are reflected to (*x*); finally, a fraction *β/*(*α* + *β*) of all dispersers die before settling, leaving only a fraction *α/*(*α* + *β*) that survive the dispersal process.

### 3.2 A single isolated patch

For the special case *m* = 1 the habitat consists of a single patch of finite length surrounded by two semi-infinite patches. This typically arises as either a single isolated patch of good habitat surrounded by poor habitat, or a large (i.e, infinite) good habitat fragmented by an unsuitable, or poor quality habitat represented by a finite interval. With *m* = 1, the dispersal kernel has 9 pieces, and *T*^−1^*W* is a 6 × 6 matrix. As with the previous case, the middle two rows and columns of *T*^−1^*W* after subtracting the identity from the upper right block are zeros, leaving 16 nonzero entries.

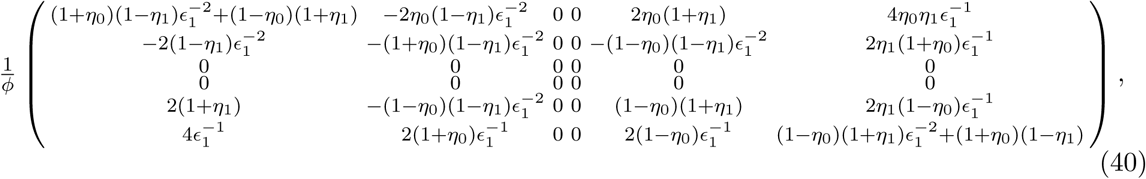

where

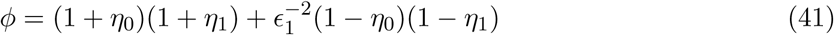

Recalling that 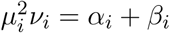, the first few pieces of the kernel are as follows:

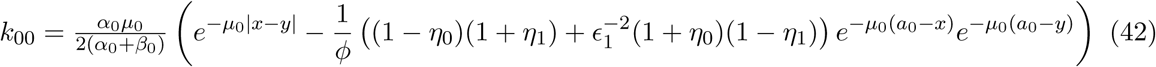

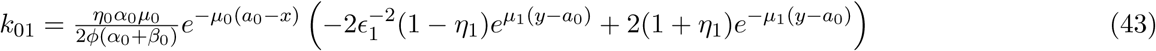

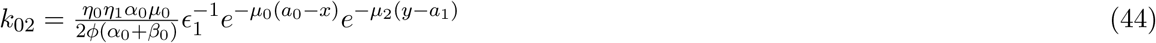

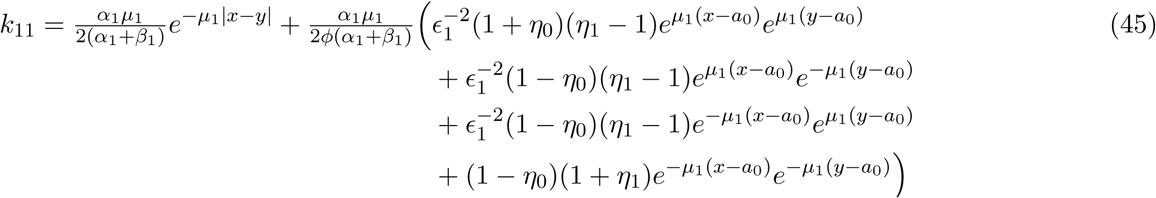

The first piece of the kernel can be expressed in terms of the homogeneous Laplace kernel and distances between the settling point, *x*, and the reflections *y*_0_ and *y*_1_ of the starting point, *y*, across the interfaces *a*_0_ and *a*_1_ respectively.

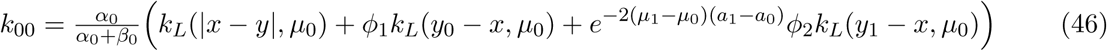

with *ϕ*_1_ = −(1 − *η*_0_)(1 + *η*_1_)*ϕ*^−1^ and *ϕ*_2_ = (1 + *η*_0_)(1 − *η*_1_)*ϕ*^−1^. It can be shown that |*ϕ*_1_| *<* 1 and |*ϕ*_2_| *<* 1, suggesting that they are fractions of dispersers reflected back into region Ω_0_ by each interface.

There are several interesting special cases appearing in the liturature where growth is zero outside the central patche and the IDE model only invovles *k*_11_. In these cases we have the following formulae for *k*_11_.

**Case 1:** If the bias at the two interfaces is equal and opposite, *z*_0_ = −*z*_1_ = *z*, and the two outside patches are identical, *µ*_2_ = *µ*_0_, *α*_2_ = *α*_0_, and *β*_2_ = *β*_0_, then 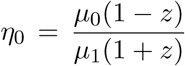, and 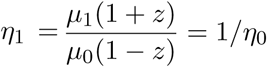. Hence,

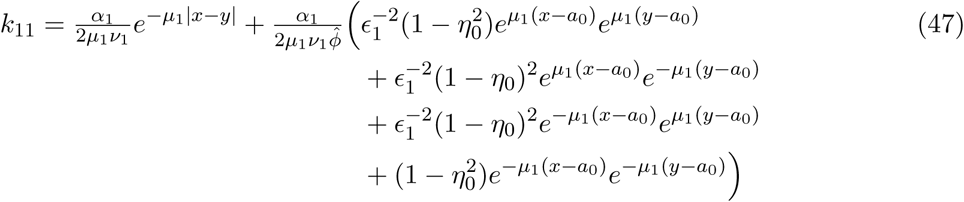

where

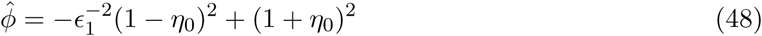

**Case 2:** If the bias at the two interfaces is equal and opposite, *z*_0_ = −*z*_1_ = *z*, and dispersal within each patch is the same, *µ*_2_ = *µ*_1_ = *µ*_0_, *α*_2_ = *α*_1_ = *α*_0_, and *β*_2_ = *β*_1_ = *β*_0_, then

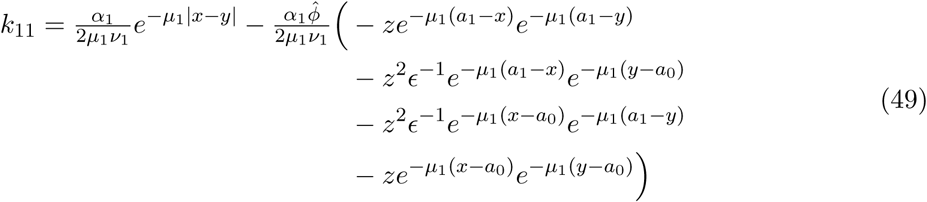

where

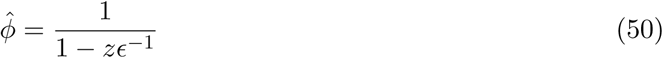

**Case 3:** If there is perfect relection back in to the central patch, *z*_0_ = −*z*_1_ = 1, then *η*_0_ → 0, *η*_1_ → ∞, and

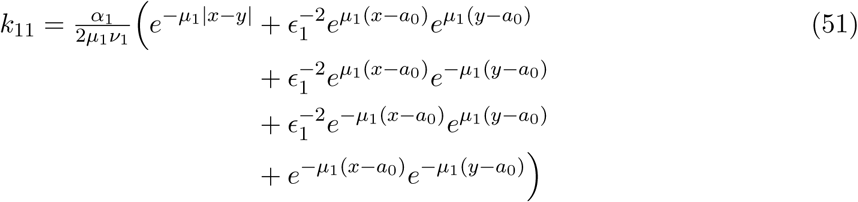

**Case 4:** If the bias at the boundaries perfectly balances the jump in kernel decay rates across the patch interfaces, *η*_0_ = *η*_1_ = 1, then *k*_11_ reduces to the truncated Laplace distribution.

which gives us

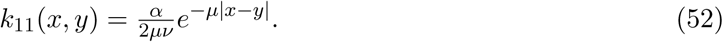

**Case 5:** Finally, we consider a case where the death rates in the exterior patches Ω_0_ and Ω_2_ are high, *β*_2_, *β*_0_ → ∞ implying *µ*_2_, *µ*_0_ → ∞ so that *η*_0_ → ∞, and *η*_1_ → 0.

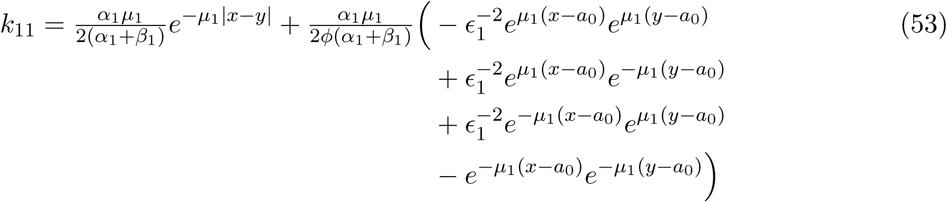

with 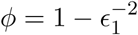. Further, as expected, the other pieces of the kernel tend to zero as all indiduals leaving the central patch die before settling.

## 4 Discussion

In this study, we assumed that the real line, as a one-dimensional landscape, is partitioned into different pieces such that there are non-equal probabilities for individuals at boundaries to move to the right or left side of the boundary. During the dispersal phase, animals may respond to habitat quality and habitat edges and these responses may affect their distribution between habitats [Allema et al., 2014]. Ecologists who have incorporated detailed and realistic behaviour into the movement process have accomplished better fits to movement data [Schick et al., 2008]. This discussion will benefit from detailed studies on movement behaviour in combination with a study on the population outcome of this behaviour. We questioned if there is a straightforward method to formulate the dispersal kernel, that came from the random walk theory, governing the probability of successful dispersal.

The *m*-patch Laplace kernel is a piecewise exponential function of two variables, *x* and *y*. Each piece, *k*_*ij*_, is a linear combination of four exponential functions (29) with coefficients representing the measure of retention at the boundaries. Moreover, *k*_*ij*_ can be expressed in terms of the distance between the settling point and the reflections of the starting point, (36). We have investigated a method to formulate the kernel with only one matrix of coefficients, *T*, independent of the location *y* but dependent on the dispersal parameters in every patch through the two parameter groups *η* and *ϵ*. With this achievement, the process of studying the effect of changing patch parameters on the model is simplified. For example, when the degree of bias at the origin increases in (36), we see *k*_10_ increases and *k*_01_ decreases. As another example, for *m* = 1, setting *η*_0_ = *η*_1_ = 1 can be viewed as balancing the bias with the ratio of the kernel decay rates across the patch interfaces.

This matrix form of the kernel will be useful for numerical simulations of IDEs, where it is necessary to integrate the product of the kernel and a reproduction function. Since the integration is over *y*, and several terms of the kernel, such as 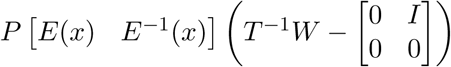, are independent of *y*. Only one part of the kernel is involved in the computation. That is, we need only integrate the product of the *j*-th column of 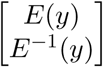 and the growth function over each Ω_*j*_. Therefore, partitioning the whole real line into *m* patches does not complicate the simulation of an IDE.

